# A data-driven approach for predicting the impact of drugs on the human microbiome

**DOI:** 10.1101/2022.10.08.510500

**Authors:** Yadid M. Algavi, Elhanan Borenstein

## Abstract

Many medications can negatively impact the bacteria residing in our gut, depleting beneficial species and causing adverse effects. To determine individualized response to pharmaceutical treatment, a comprehensive understanding of the impact of various drugs on the gut microbiome is needed, yet, to date, experimentally challenging to obtain. Towards this end, we developed a data-driven approach, integrating information about the chemical properties of each drug and the genomic content of each microbe, to systematically predicts drug-microbiome interactions. We show that this framework successfully predicts outcomes of *in-vitro* pairwise drug-microbe experiments, as well as drug-induced microbiome dysbiosis in both animal models and clinical trials. Applying this methodology, we systematically map all interactions between pharmaceuticals and bacteria and demonstrate that medications’ anti-microbial properties are tightly linked to their adverse effects. This computational framework has the potential to unlock the development of personalized medicine and microbiome-based therapeutic approaches, improving outcomes and minimizing side effects.

## Introduction

Our gastrointestinal tract harbors a flourishing and diverse community of microorganisms, collectively known as the human gut microbiome. Over the past decade, we came to appreciate how this microbiome governs individualized response to diet and susceptibility to a wide array of diseases such as diabetes and cancer^1^. Importantly, however, microbiome research has also revealed a complex and bidirectional interactions between the microbiome and numerous pharmaceuticals. On the one hand, many gut-dwelling microbes metabolize drugs, potentially affecting their toxicological, pharmacokinetic, and pharmacodynamic properties^2–6^. On the other hand, many small-molecule drugs alter the taxonomic composition of the microbiome and potentially give rise to various gastrointestinal side effects. Indeed, population-wide case-control studies in the UK and Netherlands have identified many commonly used drugs, including atypical antipsychotics, NSAIDs, and statins, as influential modulators of the intestinal microbiota^7–9^. Additionally, for specific medications, longitudinal clinical studies uncovered temporal variation following drug administration. For example, metformin – an oral glucose-lowering drug used to treat type 2 diabetes – was demonstrated to shift the microbial population in the gut, increasing the prevalence of beneficial short-chain fatty acid-producing species^10^. At the same time, metformin was also shown to increase the abundance of virulent *E. coli* strains that can cause diarrhea, bloating, and nausea – frequent adverse effects in metformin-treated patients. While such clinical studies provide some perspective on drug effects, they cannot be performed on a large number of drugs.

As an alternative approach, a complementary *in-vitro* methodology can be used to explore the potential impact of non-antibiotic drugs on gut microbes. Maier *et al*.^11^, for example, conducted a high throughput screen of more than 1,000 common drugs against 40 representative gut bacteria under anaerobic conditions. They measured the growth of each species optically over time and showed that 24% of the drugs with human targets inhibit the growth of at least one species. Similarly, others have tested 43 compounds against five different microbial communities and quantified, using mass spectrometry, the absolute bacterial abundance and proteome alterations following drug exposure^12^. This new understanding of how drugs impact the microbiome offers a novel way to improve pharmaceutical treatment as well as minimize side effects^14^. Although such recent studies have cast light on many interactions of interest, a comprehensive and complete understanding of microbiome-drug interactions is still lacking, and the incorporation of *pharmacomicrobiomics* into clinical practice is accordingly yet out of reach^16^.

To address this challenge, we integrate chemical and microbiological knowledge with a computational, data-driven, systems approach, aiming to predict the impact of a large set of drugs on the growth of microbiome members. Such computational approaches have been successfully applied to predict other types microbiome-drug relationships (*e.g*., drug biotransformation^15^), and could thus be similarly effective in facilitating large-scale characterization of drug-microbiome interactions. To demonstrate its conceptually broad applicability, we show that this approach successfully identifies the effect of pharmaceuticals on microbial strains *in-vitro* and can be further applied to predict drug-induced community changes in longitudinal animal models and clinical studies. Moreover, this methodology allows us to systematically map the influence of thousands of drugs on numerous microbial strains, uncover drug and microbial properties that underlie anti-commensal activity, and explore the connection between drug-induced microbiome compositional changes and side effects.

## Results

### Development of a machine learning model for predicting the impact of drugs on the microbiome in-vitro

We first set out to develop a random forest model that can predict the influence of any drug on any microbial species. Our model takes as input two vectors of features: one representing a specific microbial taxa and the other representing a selected drug - and aims to predict the impact of that drug on that microbe. Specifically, we characterize each microbe by the set of biochemical pathways its genome encodes and each drug by its physical-chemical properties. Overall, our model uses 148 microbial features and 92 drug features. The complete set of features used can be found in Supplementary Table 1. The model then aims to predict a continuous numerical value between 0 and 1 (that we will term throughout the paper as “impact score”), which describes the likelihood that the drug causes growth inhibition (Figure 1A; see Methods for complete details).

**Figure 1.**
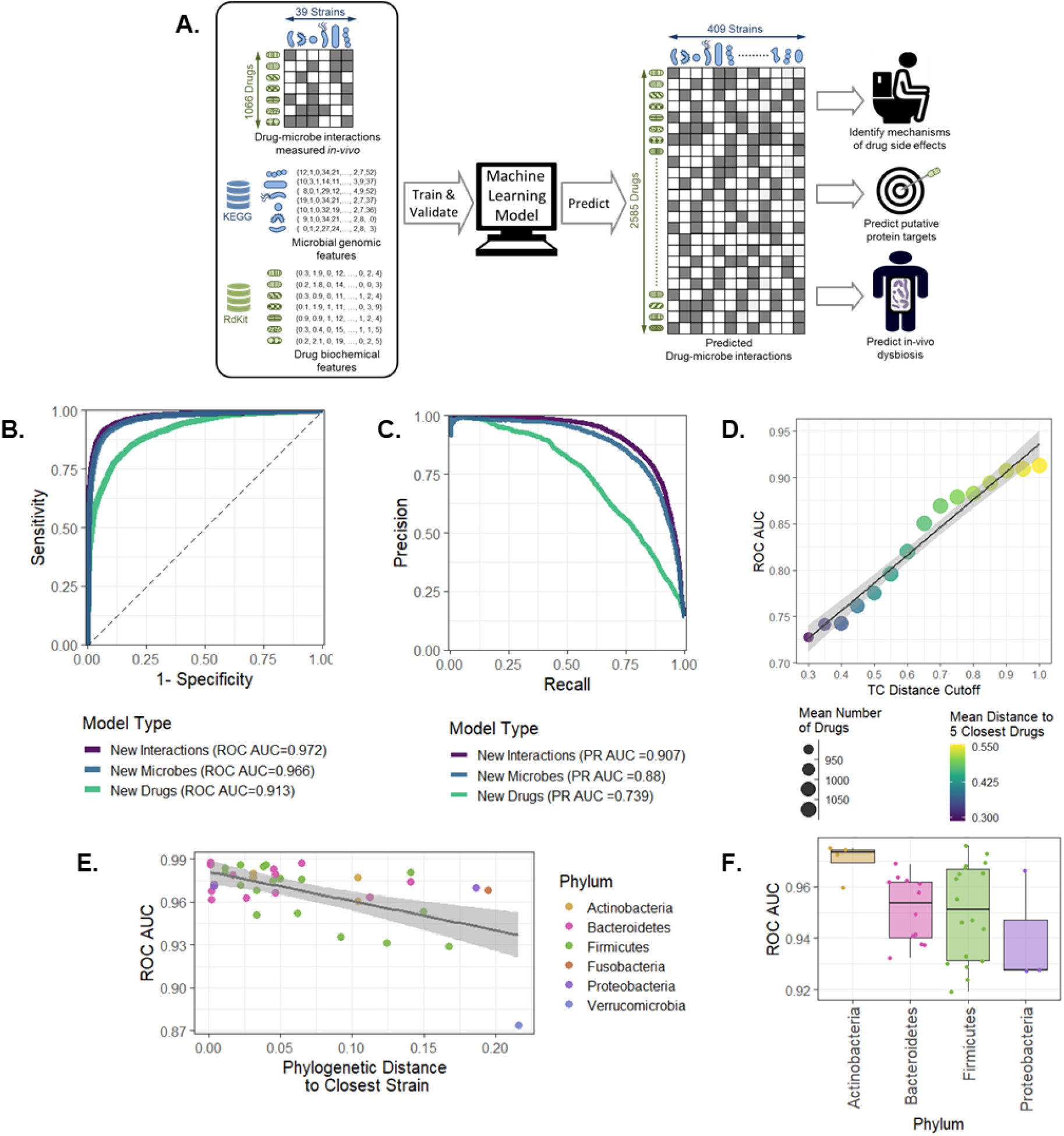
Machine learning model prediction of *in-vitro* drug-microbe interactions. **A** – A scheme of our computational framework. **B** – receiver operating characteristic (ROC) curve for in three learning settings: new drug-microbe interactions, new microbes, and new drugs. **C** – Precision recall (PR) curve for the three learning settings as in Panel B. **D** – ROC AUC score for new drug prediction as a function of the TC distance cutoff. The color scale indicates the mean distance to the five most similar drugs and the size of the circle indicates the mean number of drugs included in each cross-validation training set. **E** – ROC AUC scores for the leave-one-microbe-out model as a function of phylogenetic distance (Pearson correlation −0.574, p< 5*10^−4^). The color indicates the strain’s phyla. **F** – Decrease in ROC AUC score when all strains for the same phylum are removed from the training set. The phyla Verrucomicrobia and Fusobacteria were discarded from this analysis as each contains only a single strain.

Given this representation, we then trained our model using a large-scale dataset describing a set of in-vitro experiments where 40 microbial strains were each exposed to an array of 1,197 drugs, determining whether each drug inhibits the growth of each microbe (“0” – no effect; “1” – growth inhibition; see Methods)^11^. We tested our approach using 10-fold cross-validation across this dataset of 41,519 drug-microbe interactions. The model demonstrated excellent predictive performance in predicting new drug-microbe interactions in-vitro with an area-under-the-receiver-operator curve (ROC AUC) of 0.972 (Figure 1B). Due to the class imbalance in the dataset (ratio of 1:6.1, in favor of “0” interactions), we further report an area-under-the-precision-recall curve (PR AUC) of 0.907, indicating that the model correctly captures both types of interactions (Figure 1C). To confirm the robustness of our approach, we extensively examined other machine learning models, validated that the results cannot be attributed to statistical noise or artifacts in the data, and systematically benchmarked the model against a naive null model (see Supplementary Text 1).

Since often the impact of a given drug is relatively consistent (i.e., it either impacts most microbes or does not impact most microbes), and hence predicting a specific drug-microbe interaction when the impact of that same drug on other microbes has been used for training may not be challenging, next, we set out to examine the model’s predictive power on new drugs using a leave-one-drug-out approach. We found that while the model performance was slightly lower in these settings, it was still able to successfully predict the impact of new drugs on various microbial strains (ROC AUC of 0.913 and PR AUC of 0.739; Figure 1B, C). Notably, even after excluding antibiotics and other non-human targeted compounds, the model can still distinguish between human targeted drugs with antimicrobial activity and those without (ROC AUC 0.86), suggesting that it does not merely distinguish antibiotics vs. non-antibiotics compounds. Furthermore, to confirm that our predictions are not based solely on identifying chemically similar drugs (which accordingly have similar bioactivity), we evaluate the degree of molecular similarity between each pair of drugs using Tanimoto coefficients (TC)^17^. TC quantifies the similarity between pairs of compounds in the range of 0 (low similarity) to 1 (high similarity) based on their molecular structure. Across the 1,066 tested drugs in our dataset, the mean TC was 0.20 while the mean TC distance to the most similar compounds was 0.671. To better estimate the influence of molecular similarity on the model performance, we repeated the leave-one-drug-out approach above while excluding all compounds that are similar to the predicted drug using several similarity thresholds (Figure 1D). We confirmed that even when all compounds with a TC distance of 0.3 or higher are removed from the training set, the model retains reasonable predictive power with ROC ACU of 0.73 (with the performance further improving as more structurally similar compounds are included in the training set).

Further, using an analogous leave-one-microbe-out approach, we inspected the model capabilities in predicting the response of new microbial strains. The model demonstrates significant predictive power with a ROC AUC of 0.966 and PR AUC of 0.88 (Figure 1B, C). Examining the prediction accuracy of our model for strains of various phyla, we further confirm accurate predictions for the two main phyla of the human gut microbiome, Bacteroidetes and Firmicutes (ROC AUC 0.974, 0.966, PR AUC 0.91, 0.899, respectively). As expected, the model performance improves when phylogenetically similar strains are included in the training set, indicating that phylogenetic similarity translates to related drug-response (Figure 1E). To further determine the robustness of the model, drug response was predicted separately for each microbe when excluding *all* other microbes from the same phylum from the training set. Even under these conditions, performance remains robust with all strains achieving a ROC AUC above 0.92 (Figure 1F).

Lastly, to further validate our model’s predictions on independent datasets, we evaluated its performances on additional in-vitro screening results. First, we utilized data describing 25% inhibitory concentration values (IC25) for 25 human-targeted medications against selected microbial strains^11^. For each drug, we compared our model prediction for that drug’s impact on each strain against the measured IC25. Of the 25 drugs tested, we found a negative correlation between the strain’s impact score and the measured IC25 values in 21 compounds, with FDR corrected p<0.2 for 10 drugs (Spearman correlation test; Supplementary Figure S1A). Second, we used an independent dataset describing the impact of 43 drugs on *ex-vivo* human fecal samples^12^. Evaluating the prediction accuracy of our trained model on this new validation dataset, we found that it exhibits good transferability in predicting specific drug-microbe interactions, with a ROC AUC of 0.70 (Supplementary Figure S1B). We also estimated the predictions on new drugs and new microbes using a leave-one-out approach as before, finding only a minor loss in performance (2-3% decrease in ROC AUC; Figure S1B). These findings confirm that the model successfully predicts drug-microbe interactions on various microbial strains, even when applied to datasets obtained from different experimental setup.

**Supplementary figure S1.**
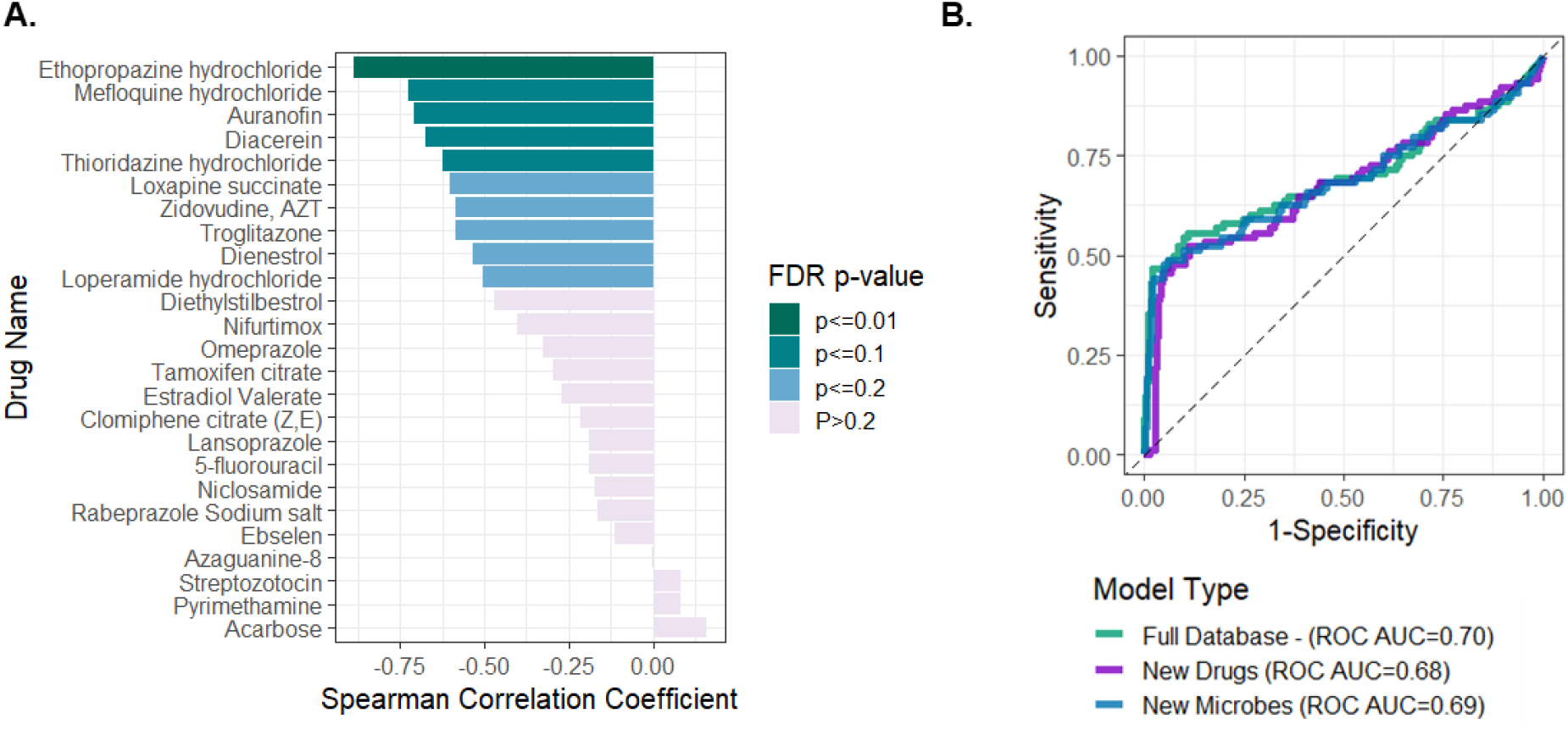
Validation against additional in-vitro datasets A. Correlation between the models predicted impact scores and *in-vitro* determined IC25 measurements. Experiments were conducted on a range of microbial strains (16 drugs against 12 microbes, 6 drugs against 16 microbes, and 3 drugs against 25 microbes). Each panel illustrates the results for one of these 25 drugs. B. Receiver operating characteristic (ROC) curve for independent predictions of 43 drugs against 19 microbes in three prediction settings: full dataset, new microbes, and new drugs.

### Structural and microbial features that influence drug impact on the microbiome

We next examined the features that contributed most to our model’s predictions to further reveal intriguing and valuable insights into the factors that impact drug-microbe interactions. To this end we used a permutation importance method to calculate statistical significance and contribution of each feature^18^. We first examined which drug features are most informative for predicting antimicrobial activity. Importantly, we found that measures of compound lipophilicity (MolLogP) and charge distribution (PEOE) are the most significant contributors for predicting antimicrobial activity (p < 0.05, Figure 3A). Surface charge properties (such as Total Polar Surface Area - TPSA), hydrogen bonding, and topological features (such as Kappa and Chi connectivity indices) were also found to be contributing. These features are consistent with known properties of antimicrobial compounds^19^. Likewise, topological drug descriptors were also found to be informative, and indeed such features are known to be relevant for antibiotic accumulation inside bacteria^19,20^. Additionally, we examined whether the relative importance of drug features varies between different microbes, training the model on one strain at a time, calculating feature significance for each such model, and using a principal component analysis (PCA) to explore variation in these feature importance sets. We found that strain-specific drug feature importance scores cluster according to both phylum and gram stain (Supplementary Figure S2A,B; PERMANOVA p<0.01 and p<0.05 for phylum and gram stain, respectively). Interestingly though, the main difference in feature importance scores between microbial strains is observed in the topological, charge, and lipophilicity descriptors (Supplementary Figure S2C), while kappa1, molecular weight, and number of hydrogen bonding acceptors further differentiate between gram-positive and gram-negative strains (p < 0.05; t-test). Indeed, hydrogen bonding and size have been reported to differ between gram-positive and negative antibacterial compounds^19^. We also examined the feature importance obtained for a model trained only on human targeted drugs as these drugs, unlike antibiotics, were not optimized for cellular penetrance. Compression between the two lists of important drug features revealed increased importance of topological features in human targeted drugs (t-test, p< 0.05), with a general positive correlation between the two lists (Pearson correlation 0.523, p< 5*10^−6^).

**Figure 2.**
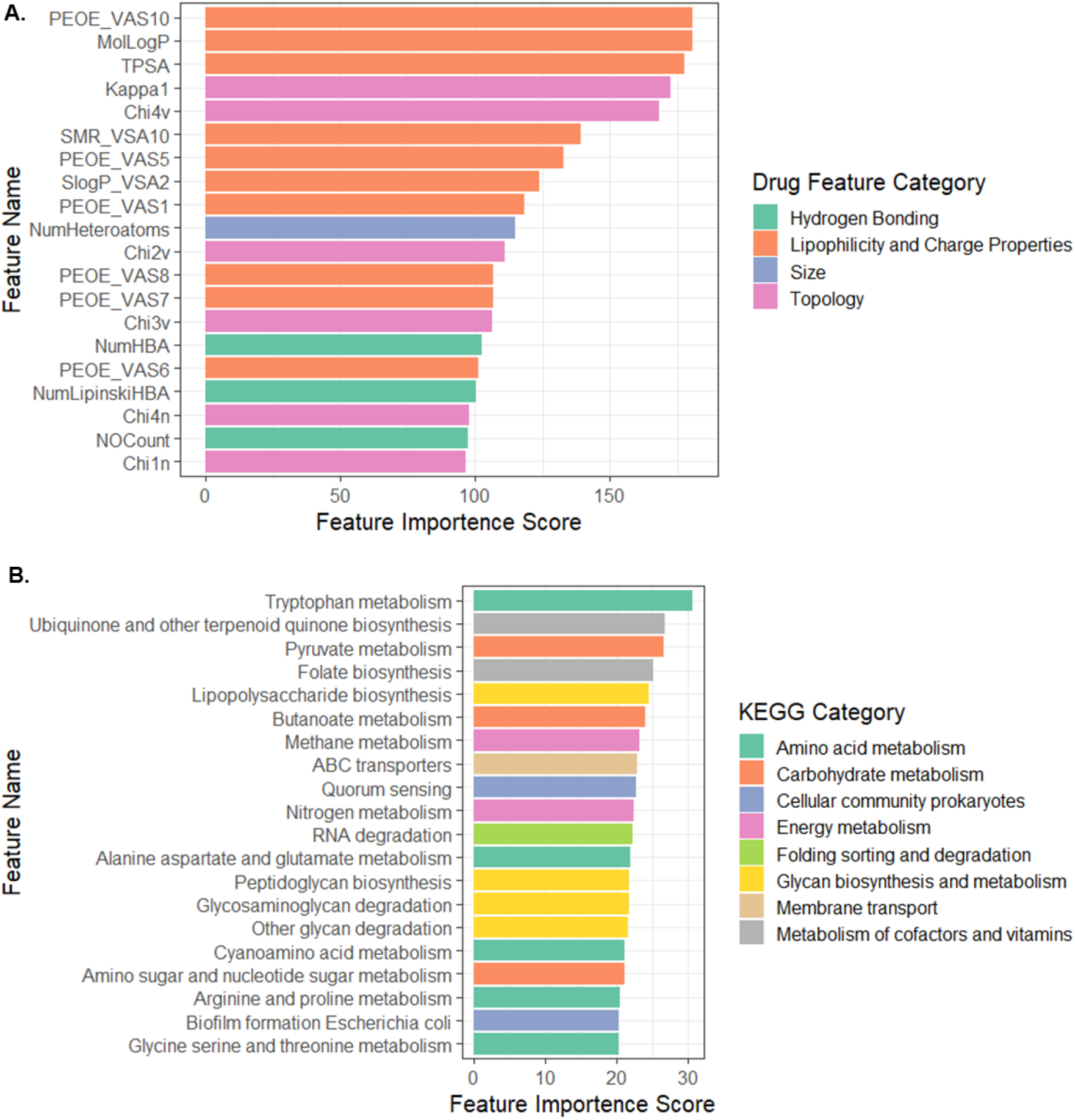
The model’s drug and microbial feature importance scores. **A** – The top 20 drug features with highest feature importance score colored according to the drug feature category. **B** – The top 20 microbial features with highest feature importance score colored according to KEGG categories. All features included in these plots are statistically significant (p <0.05).

**Figure 3.**
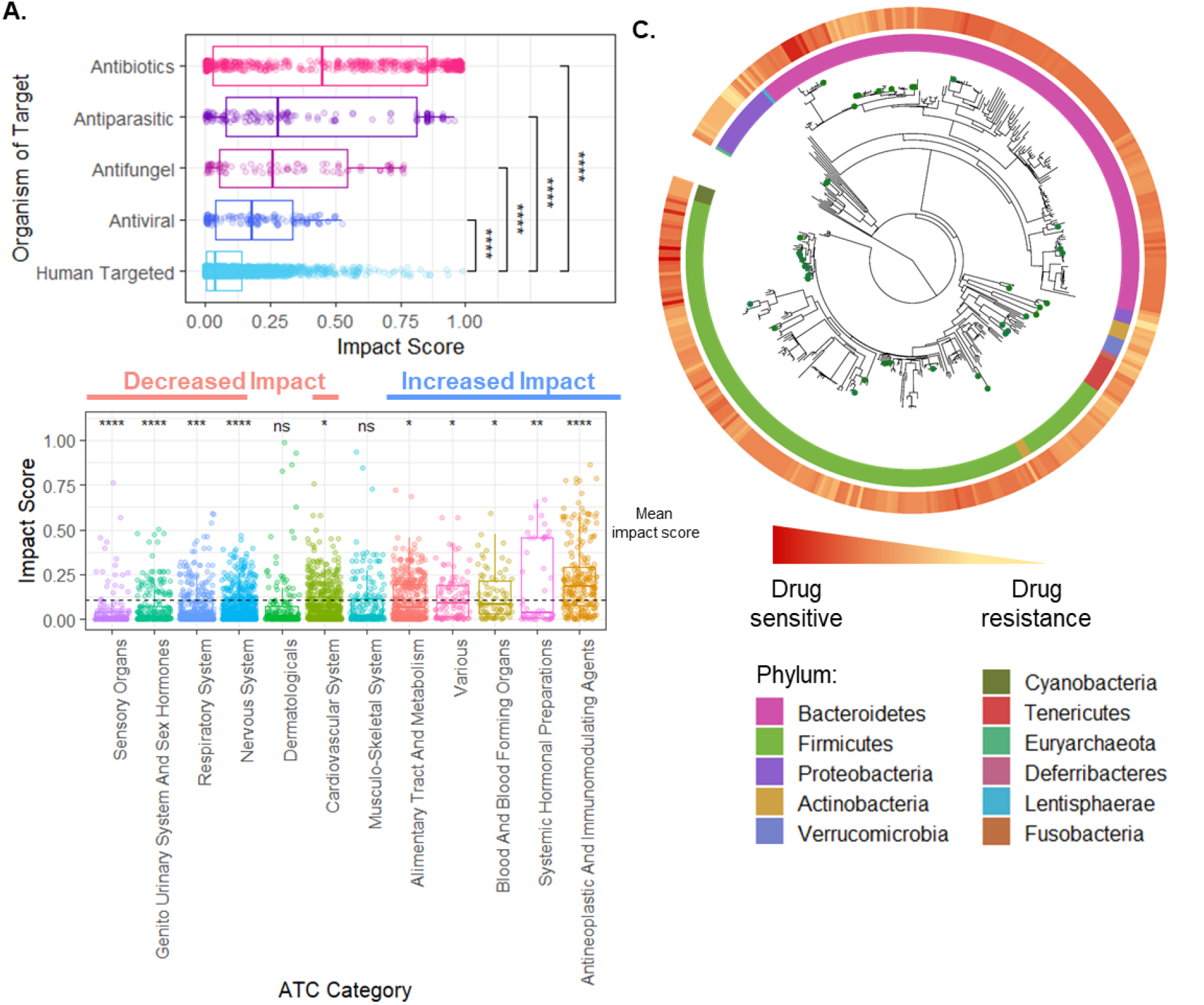
The landscape of drug impact on microbiome members. **A** – Box plots describing the difference in impact scores between human target drugs and drugs targeting various microorganisms. Significance was calculated using the Tukey’s test. **B** – Box plot describing the difference in impact scores between human targeted drugs according to Anatomical Therapeutic Chemical (ATC) classifications. The symbols above each label indicate the statistical significance in compression with the mean impact score (t-test, ANOVA test further show that the differences between ATC categories are statistically significant (p < 1*10^−15^). **C** – Drug impact across the tree of life. Phylogenetic tree for 409 microbiome members constructed based on their 16s rRNA gene. The outer circle denotes the drug sensitivity index, and the inner circle denotes the phylum. Taxa marked with green dots are those included in the *in-vitro* screen (n=39). ns – not significant; * p < 0.05; ** p < 0.01; *** p < 0.001; **** p <0.0001

We further investigated in a similar manner which microbial feature contributed most to the prediction of drug-microbe interactions. Interestingly, we found that 54 biochemical pathways from 16 KEGG categories significantly contributed to the model (p <0.05, Figure 2B). Significant microbial features include known cellular processes that are essential for antibiotic resistance. For example, indole – a byproduct of tryptophan metabolism, the top-ranking feature in this list, is known to cause antibiotic tolerance in bacteria^21^. Similarly, co-factor biosynthesis pathways such as ubiquinone production and folate biosynthesis are known to regulate oxidative stress^22,23^. Not surprisingly, features that encode for membranal structure and transport, lipopolysaccharide biosynthesis, and ABC transporters were also found to have high importance^24^. We further found that the microbial features that contribute to the original model and those that contribute to a model trained on only human targeted drugs are highly correlated (Pearson correlation 0.904, p<5*10^−16^), suggesting that similar genomic components might be utilized for resistance against antibiotics and nonantibiotic drugs.

**Supplementary figure S2.**
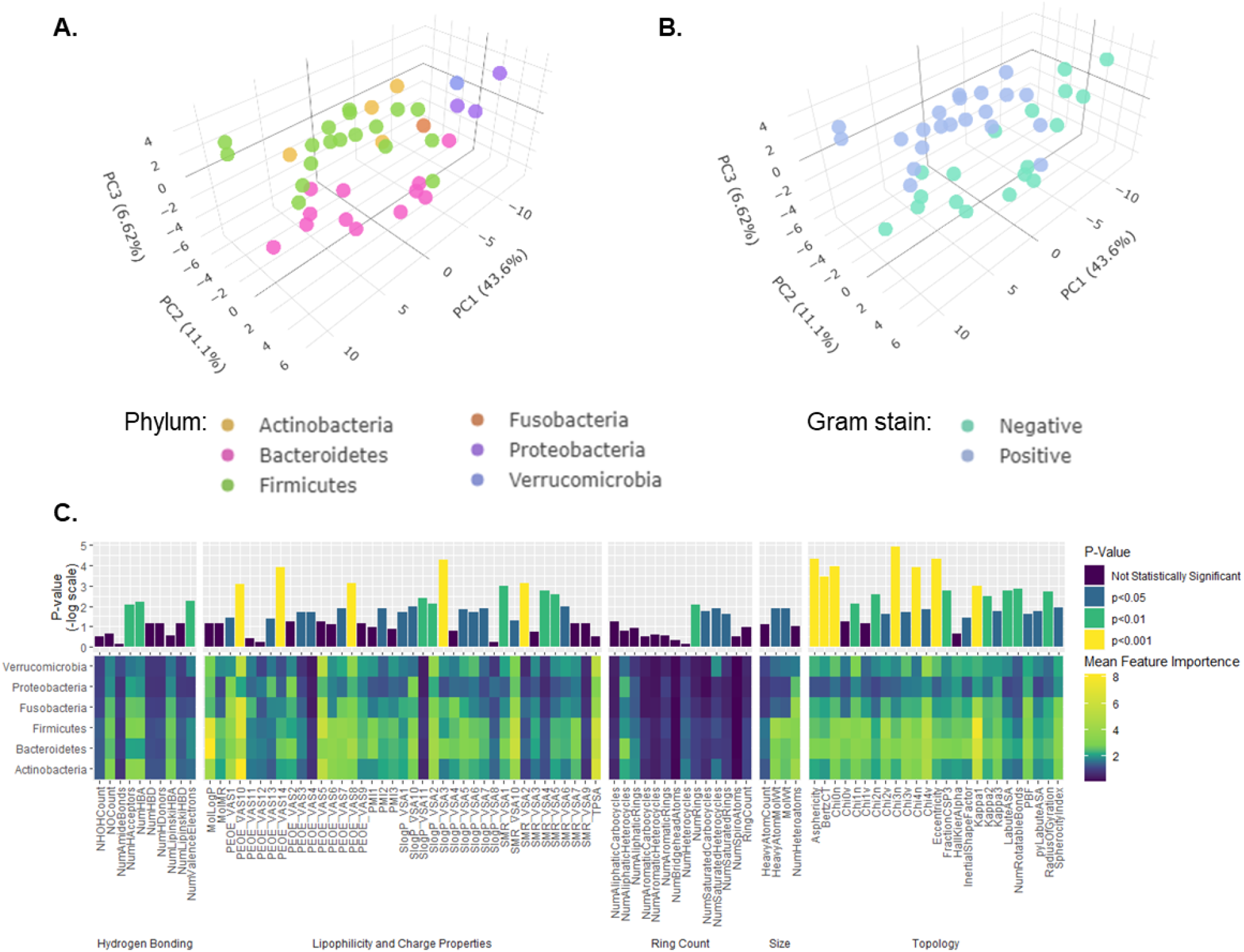
Differences in the importance of drug features between different microbial taxa. Principal Component Analysis (PCA) of the importance of drug features. Each point represents a single microbial strain and is clustered according to **A** – Phyla (PERMANOVA, p = 0.001) and, **B** – Gram stein (PERMANOVA, p = 0.017). **C** – Differences in mean feature importance between phyla. Top panel – Two-way ANOVA FDR corrected p-value for each feature. Bottom panel – Mean feature importance for each phylum.

### The landscape of drug impact on the human microbiome

Having established and benchmarked our model using available data on drug-microbe interactions *in-vitro*, we set out to explore the complete landscape of interactions between drugs and members of the human gut microbiota. Toward this aim, we collected *all* clinically approved small molecule drugs from the DrugBank database^25^ and calculated their physio-chemical properties. Similarly, to represent the diversity of the intestinal bacterial community, we used metagenomic data from a representative, healthy western population^26^ (see Methods for details). We then trained our model on the complete *in-vitro* data described previously and systematically predicted the impact of each of 2,585 drugs on 409 human microbiota members, resulting in a rich catalog of 1,057,265 drug-microbe interactions (Supplementary Table 2). Notably, more than 62% of the drugs and 90% of the microbial taxa have not been tested *in-vitro* before.

Using this catalog, we first sought to identify patterns in the drug-microbe interaction landscape, highlight drug- and microbe-specific determinants of such interactions. Examining the predicted impact of each drug on the various taxa revealed that, perhaps not surprisingly, antibiotics cause the most impact on the microbiome. Likewise, pharmaceuticals whose targets are non-human, such as antivirals, antifungals, and antiparasitic drugs, also impact many species (Figure 3A). These compounds were designed to penetrate through the cellular membrane and disrupt vital biochemical pathways that could be shared among organisms. Inspecting human-targeted drugs, we found distinct differences in their impact on microbiome species based on the anatomical system they target (Figure 3B, two-way ANOVA, p<1*10^−15^). Drugs with antineoplastic activity show the highest lethality, followed by drugs that act on the hormonal system, blood-forming organs, and the alimentary tract. In contrast, medications affecting the sensory, genitourinary, and respiratory systems show the lowest average activity (Figure 3B). Considering specifically the top 20 drugs (in term of their mean impact on microbial taxa) that have not been screened in the original database, we found several compounds with reported *in-vivo* anti-commensal activity such as, the immunosuppressants Sirolimus^29^ and Tacrolimus^30^ and the antineoplastic agents’ Somatostatin^31^ and Eribulin^32^.

Next, we used the above catalog to examine differences in drug sensitivity (which we define as mean impact score across all compounds) across the microbial tree of life (Figure 3C). We found that microbiome members from the Verrucomicrobia and Proteobacteria phyla showed generally higher resistance to drugs. This resistance could be attributed to their rich capacity for drug metabolism and the low permeability of double-layer membranes^33^. Comparing the sensitivity of the two main phyla of the human microbiome, Firmicutes and Bacteroidetes, whose ratio is known to be associated with multiple lifestyles and clinical factors^34^, has also exhibited intriguing patterns. Specifically, while both phyla are predicted to be drug-sensitive, over 80% of the drugs have some specificity for one or the other (t-test, FDR p<0.05). While antiparasitic, respiratory, and nerve system drugs have a more specific impact on Bacteroidetes, hormonal preparations are more specific toward Firmicutes (t-test, FDR p<0.05).

Lastly, we aimed to identify a possible mechanistic explanation for the anti-commensal properties of human targeted drugs. Unlike antibiotics, most human-targeted medications do not have a recognized microbial target. For this purpose, we obtained manually curated protein target information from the DrugBank database and mapped these proteins to their microbial orthologs (see Methods). We found 899 drug-microbial target interactions between 497 compounds and 198 proteins. We then searched for drugs that had significantly different impact scores between taxa with and without the target protein, identifying 201 such pairs (Wilcoxon rank-sum test, FDR p < 0.05, Supplementary Table 3). Among these are compounds with previously identified targets, such as nucleotide analogs which inhibits thymidylate biosynthesis, a proposed target for antibiotics^35^, and streptozotocin, which inhibits intracellular protein glycosylation^36^. We further identified possible targets of compounds with characterized antimicrobial activity but without recognized microbial protein targets including, sevoflurane inhibition of calcium-transporting ATPase^36^ and sitagliptin inhibition of Dipeptidyl peptidase 4 orthologues^37^ (Supplementary Figure S3). This analysis could pave the way for deeper understanding and possible prevention of this unwanted off-target activity.

**Supplementary figure S3.**
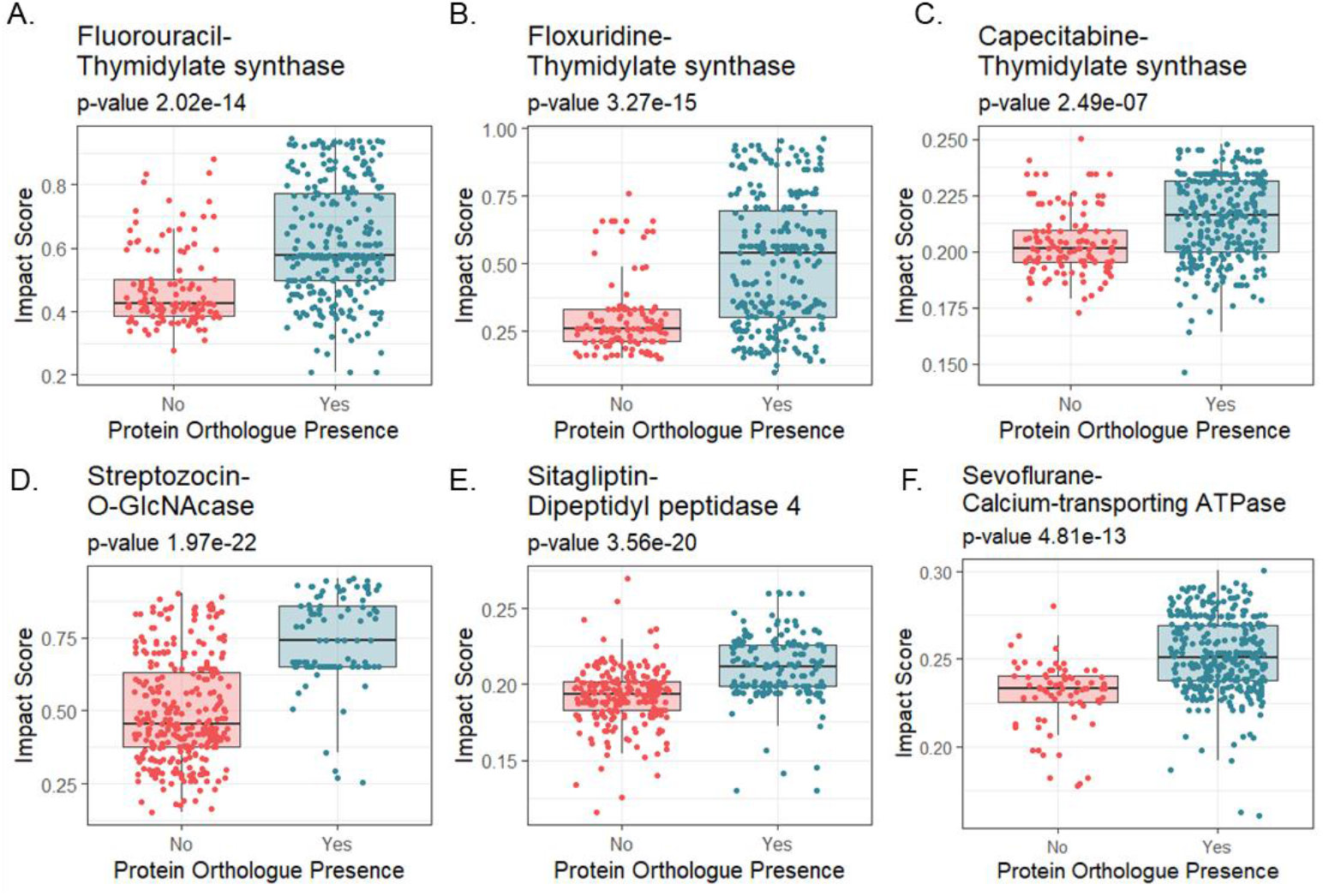
Differences in impact score between taxa according to the presence of protein target orthologues. **A-F** – Difference in impact score for Fluorouracil (A), Floxuridine (B), Capecitabine (C), Streotozocin (D), Sitagliptin (E), and Sevoflurane (F) between taxa with and without orthologue to thymidylate synthase (A), thymidylate synthase (B), thymidylate synthase (C), O-GlcNAcase (D), Dipeptidyl peptidase 4 (E), and calcium-transporting ATPase (F). Significance was calculated using Wilcoxon rank-sum test and FDR corrected p-values.

### Prediction of *in-vivo* drug-induced dysbiosis

Next, we set out to evaluate the ability of our computational framework to predict the impact of various drugs on the microbiome community *in-vivo*. Extending previous validation^11^, here we aimed to test the impact of specific drugs, both on the much larger set of microbes included in our model’s predictions and in terms of observed variation in the complete microbiome composition profile following drug administration. Although the intestinal ecological dynamics are far more complex than the those of single strain *in-vitro* experiments, we hypothesized that taxa predicted to be strongly impacted by a given drug, would exhibit reduced abundances after pharmacological intervention with that drug. Towards this aim, we collected metagenomic sequences from longitudinal studies in which subjects were sampled before and after pharmaceutical treatment (Supplementary Table 4). This study design allows direct evaluation of the impact of the drug on intestinal bacteria, while minimizing other confounding factors that can arise in case-control studies such as inter-individual variability in the microbiome baseline composition. To examine our hypothesis, we compared model predictions of each taxon with the change in their relative abundance after treatment (see Methods).

First, we examined the model’s ability to predict microbiome alternation in human clinical trials. Omeprazole is a common proton pump inhibitor used in the treatment of gastroesophageal reflux disease, and is associated with decreased diversity of intestinal species and increased risk for *Clostridium difficile* infections^38^. Comparing microbiome composition before and after Omeprazole treatment with our model’s predicted impact has demonstrated that our model correctly captures the effect of this drug on the microbiome, assigning higher impact score to taxa whose relative abundances were reduced following omeprazole administration (Figure 4A). Importantly, although we trained the model on merely 39 strains, it was able to predict the impact on a microbiome community of 153 different taxa, of which only 15%were tested *in-vitro* against omeprazole while the rest represent novel predictions of the model. Moreover, repeating this analysis while separating taxa tested *in-vitro* from those that were not, we found that the statistical significance reported above is primarily driven by those members of the microbiota not yet tested (Supplementary Figure S4A). We further applied our model on data available from two pre-clinical animal models treated with Paclitaxel (also known as Taxol), a chemotherapy used to treat various solid cancers^39^, and Methotrexate, the first-line therapy for rheumatoid arthritis^40^. In both cases, our model successfully predicted the drug-induced changes in the gut microbiota (Figure 4B-C). As for Omeprazole, most of the taxa were previously unscreened *in-vitro* and are driving for statistical significance of these analyses (Supplementary Figure S4B-C). Additional analysis further confirmed that in all cases above, the results cannot be attributed merely to statistical noise since randomization of features resulted in total statistical significance loss (Methods; Supplementary Figure S4D-E).

**Figure 4.**
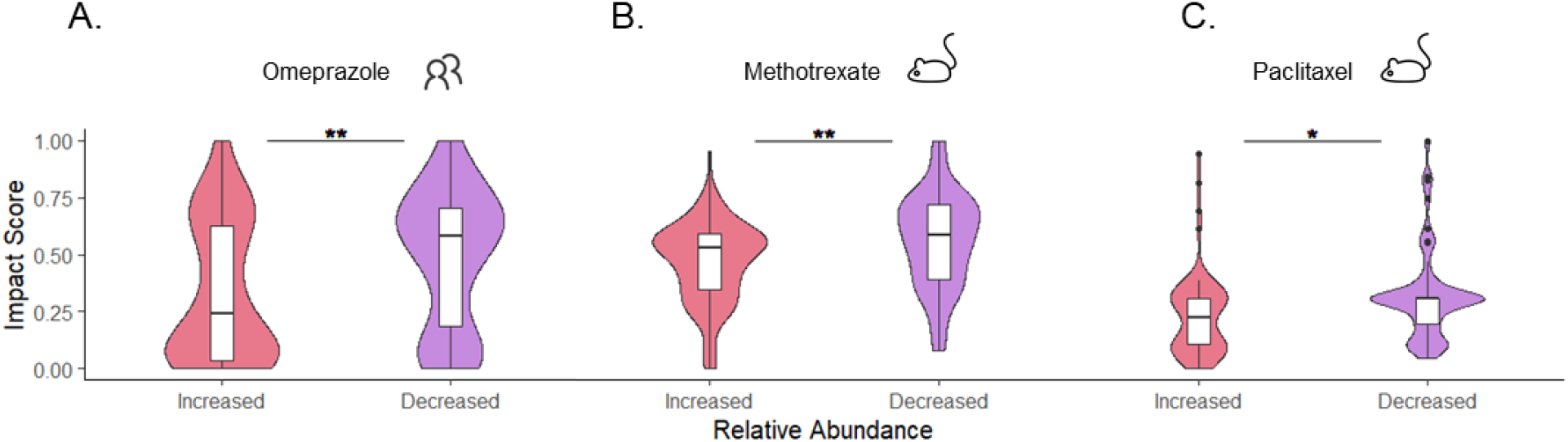
Prediction of *in-vivo* drug-induced dysbiosis. Violin plots illustrating the difference in predicted impact scores between taxa with significantly increased vs. decreased abundance following drug administration of **A** – Omeprazole (human), **B** – Paclitaxel (mouse), and **C** – Methotrexate (rat). * p < 0.05, ** p < 0.01, *** p < 0.001

**Supplementary Figure S4.**
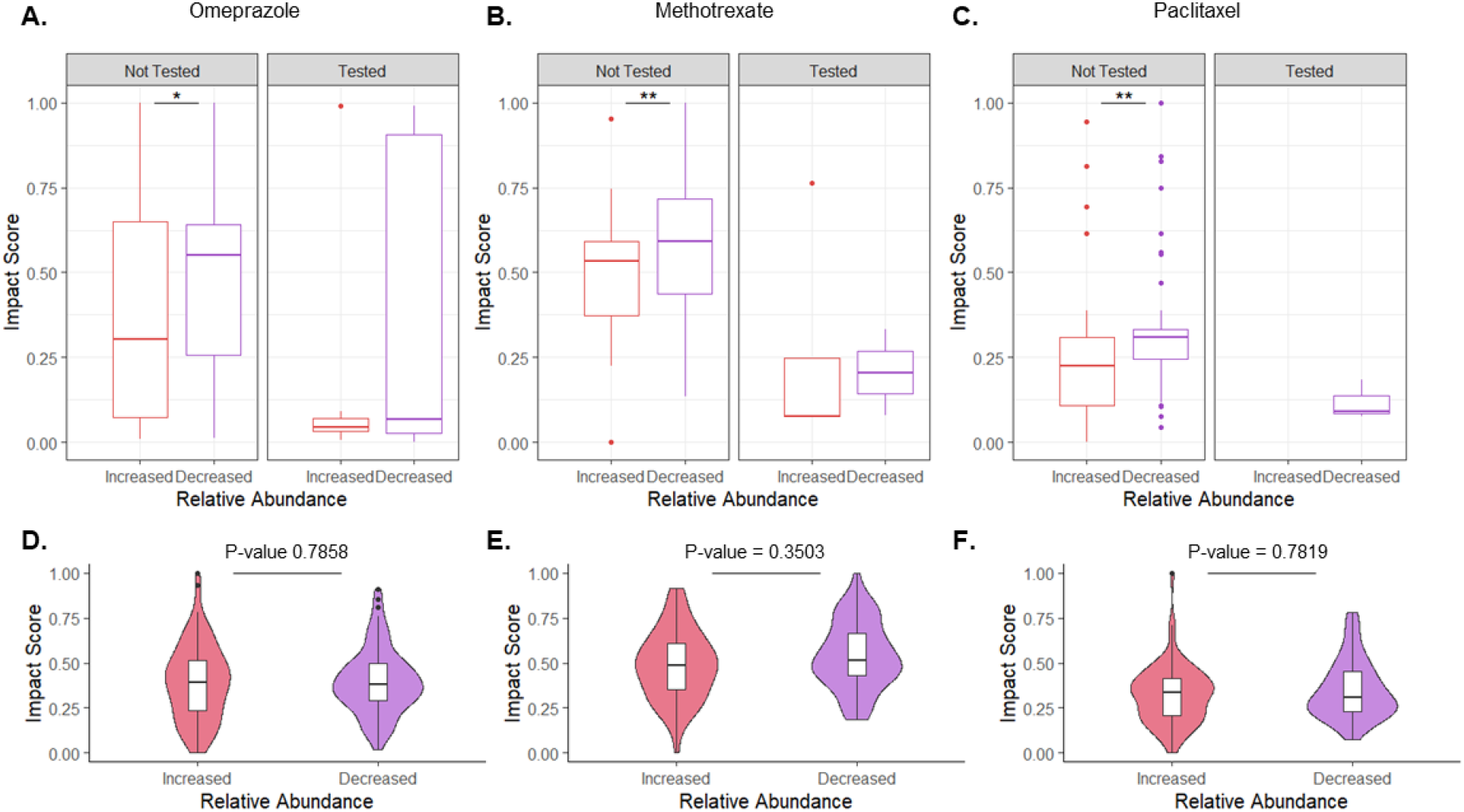
Controls for *in-vivo* model predictions. **A-C** – Compression in prediction scores between taxa with increased and decrease abundance after drug administration. Results are shown separately for taxa present and absent from the *in-vitro* dataset. **D-E** – Compression in prediction scores between taxa with increased and decrease abundance following drug administration using a model with shuffled features. Statistical significance was computed using t-test. * p < 0.05, ** p < 0.01, *** p-value < 0.001

Interestingly, this ability to successfully predict drug-microbiome interactions *in-vivo*, further allows us to predict microbiome-wide drug sensitivity across global healthy populations. For example, immigration to the United States from Thailand has been previously shown to be associated with loss of bacterial diversity and fiber degradation capacity^41^. Applying our model to predict the weighted mean impact score (as a proxy for personalized microbiome drug sensitivity) of individuals from various cohorts suggests that westernization is also associated with increased sensitivity to pharmaceutical compounds (see Methods). Specifically, our model predicts that Individuals born in the USA, regardless of whether they are of Thai descent or not, are more sensitive to drugs than individuals born in Thailand (Supplementary Figure S5). This preliminary analysis demonstrates the potential of computational prediction of drug-microbiome interactions, facilitating the generation of novel population-level hypotheses about the link between gut microbes, drugs, and impact on the host.

**Supplementary figure S5.**
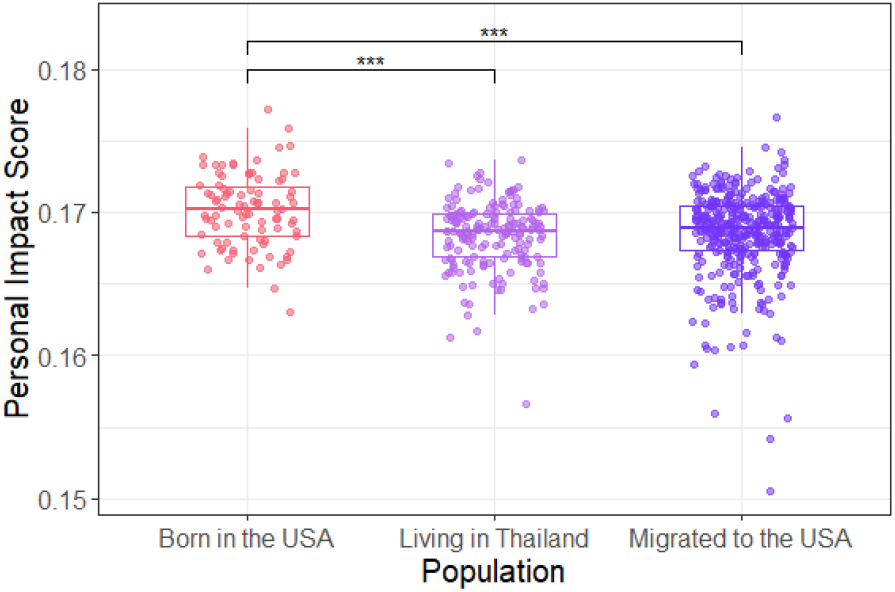
Influence of westernization on the resistance of the microbiome to drugs. The difference in mean personal impact score between three populations: Thai population in Thailand, Thai migrated to the United States (USA), and individuals born in the USA (Tukey’s honest significance test). * p < 0.05, ** p < 0.01, *** p < 0.001

### Characterizing links between adverse drug reactions and anti-commensal activity

Lastly we examined whether the anti-microbial properties of drugs may be related to their side effects. Specifically, following previous studies that highlighted a link between drug-induced dysbiosis and side effects in a handful of medications, we wondered whether using our model’s predictions we can observe this link on a much larger scale. To this aim, we collected and curated side effects reports from the Side Effect Resource (SIDER) database^42^ for all non-antibiotic small molecule compounds (n=771). Then, we predicted the impact of each of these drugs on the representative microbiome community described above and calculated the mean impact score of each drug across all taxa. Comparing the drugs’ impact scores with their cataloged side effects, we found that both gastrointestinal and infection-related adverse effects were strongly associated with the drug’s impact on the microbiome. Specifically, non-antibiotic medications with a high incidence of these side effects exhibited a significantly higher impact score compared to those with a low incidence of these side effects (Figure 5A-B), suggesting that perhaps similarly to antibiotics^11^, drugs with an extensive impact on the intestinal community might facilitate gastrointestinal side effects and colonization of pathogenic bacteria.

**Figure 5.**
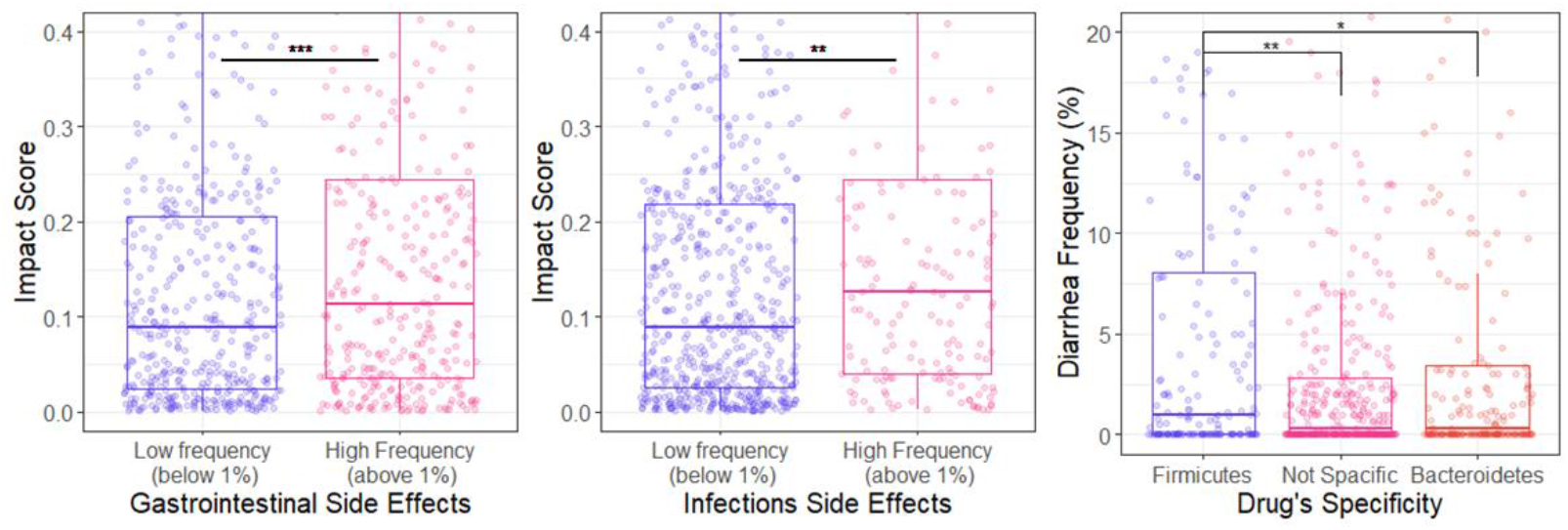
Association of anti-commensal activity and adverse drug reactions. Box plots illustrating the difference in predicted impact scores between drugs with **A** - high vs. low frequency of gastrointestinal drug adverse effects, and **B** – high vs. low frequency of infections drug adverse effects. **C** – Difference in diarrhea frequency between drugs with impact specificity to Firmicutes and to Bacteroidetes, as well as drugs without specificity to any phyla (Tukey’s honest significance test). * p < 0.05, ** p < 0.01, *** p < 0.001

Based on this finding, we next examined whether specific dysbiosis patterns might be associated with various adverse effects. Above, we found that each drug tends to affect mostly strains for one of the two main phyla of the human microbiome, Firmicutes or Bacteroidetes, with smaller effect on strains from the other. Here, we compared for each drug the difference in its impact on these two phyla (as a measure of its specificity), and calculated the association between each side effect and this drug specificity. We found that diarrhea is significantly more common in patients receiving drugs that are more harmful to Firmicutes than to Bacteroidetes, with a ~66% increase in incidence rate (ANOVA, p = 5*10^−3^, Figure 5C). Interestingly, in line with our findings, stool consistency was positively associated with Bacteroidetes:Firmicutes ratio in a healthy woman population^43^. Based on those findings, we hypothesize that the mode of drug-induced dysbiosis may explain the patterns of certain adverse effects.

## Discussion

In this study, we developed a machine learning approach that integrates chemical properties and genomic content to predict the impact of drugs on microbiome taxa. We demonstrated the utility of this approach at a range of *in-vitro* and *in-vivo* settings, from pairwise microbe-drug experiments, through animal models, to clinical trials. Importantly, beyond its ability to predict the impact of specific drugs on specific microbes, this approach uncovers the determinants behind the interactions between pharmaceuticals and microbes and systematically maps the interactions between every drug and all representative members of the human microbiome. Given this extensive large-scale catalog, we were able to further provide intriguing hypotheses about the drug sensitivity of the microbiomes of various global populations and found a strong association between the anti-commensal properties and gastrointestinal side effects.

Despite these advances, our approach is not without drawbacks and relies on several main simplifications. First, the anti-commensal effect of a given drug clearly depends on its bioavailability in the site of action, yet our model relies on data from a high-throughput screen in a fixed concentration. Although the authors demonstrated that this concentration is within an order of magnitude from the intestinal concentration for most drugs, it might be lower or higher for some compounds and could be influenced by both host and microbial factors. Further, our method broadly captures the differences between microbial taxa, possibly ignoring some aspects of strain-specific variation. Indeed, the presence of specific xenobiotic-metabolizing enzymes and multidrug resistance transporters are well known to impact antimicrobial and anti-drug resistance^44,45^. Lastly, rich ecological dynamics between microbiome community members and their interactions with epithelial and immune cells along the gastrointestinal tract could markedly alter the impact of various drugs on the microbiome^46^.

Looking forward, dissecting the plethora of interactions between drug, microbes, and the host holds great promise for clinical applications^5,14^. In the last decade, researchers have characterized the cardinal role of the human microbiome on pharmaceutical treatment, highlighting various processes such as biotransformation to inactive or even toxic compounds^47–49^, alternation of pharmacokinetics and pharmacodynamic properties^3^, and adverse reactions associated with drug-induced dysbiosis^50,51^. Excitingly, such observations are currently being translated into new clinical interventions protocols, optimizing treatment outcomes by developing inhibitors for drug-metabolizing enzymes^52–54^, and manipulating the microbial community using prebiotics and fecal microbiome transplants^55^. Our methodology could substantially complement these efforts, guiding future drug development attempts by providing crucial information about potential personal alterations to the microbiome following pharmaceutical treatment and identified possible mechanistic explanations for the anti-commensal activity and side effects. The use of this and other computational tools^15^ could accordingly benefit future efforts in pharmacological and microbiome research, paving the way for personalized pharmaceutical therapy and tailored microbial interventions.

## Data and methods

### Machine learning model, data, and evaluation

We implemented a machine learning model (see detailed below) to estimate the impact of each drug on each microbiome member. The model represents each drug-microbe pair as a vector of features. Drugs are described by a set of physical-chemical and structural properties, obtained from SMILES representation, using RdKit, an open-source chemoinformatic program^56^. Each microbe is described by its KEGG pathways’ scores based on its genomic content, calculated using a previously published method^57^. Briefly, the score of each KEGG pathway represents the number of KEGG orthology groups (KOs) in that pathway (with KOs associated with several pathways being partitioned between these pathways). The list of all drug and microbe features is available in Supplementary Table 1.

To train and validate our model, we utilized data previously published by Maier *et al.^11^* In brief, the data describes an *in-vitro* screen of 1197 compounds against 40 microbial strains. For 36 of the 40 tested microbial strains, we obtained KEGG KO annotation from the IMG GOLD database^58^. The genomes of three strains (*Ruminococcus gnavus VPI C7-9, Bacteroides fragilis enterotoxigenic 20656-2-1*, and *Ruminococcus torques VPI B2-51*), whose KO annotations were not available in IMG, were downloaded from NCBI and annotated using BlastKOALA^59^. We discarded one strain (*Clostridium perfringens C36*) from further analysis as its genome wasn’t publicly available. Out of the 1197 screened drugs, we matched 1066 compounds to the DrugBank database^25^ by mapping their IDs to stitch5 IDs and curating this mapping manually (unmatched drugs were mostly non-pharmaceuticals or veterinary drugs). A binary interaction score (“0” – no interaction; “1” – growth inhibition) was given to each microbe-drug pair according to the definition in Maier *et al*.^11^

Given these data, we trained several ML models including random forest (RF), supported vector machine (SVM) with either polynomial or radial basis function (RBF) kernels, and three regularized logistic regression models (ridge, lasso, and elastic net) with default hyperparameter. We intentionally focused on relatively simple and commonly used models to avoid the risk of overfitting. The performance of each model was estimated using 10-fold cross-validation and evaluated by ROC AUC and PR AUC. The above models were implemented in R version 4.0.2^60^, using “tidymodels” package suite^61^. Specifically, we used “ranger” for RF model^62^, “keranlab” for SVM algorithms^63^, and “glmnet” for logistic regression models^64^.

Evaluation of model performance on new microbes and new drugs was carried out using a leave-one-out cross-validation strategy. To determine how phylogenic and molecular similarities affect new microbe and drug predictions, we used phylogenic distance based on the 16S rRNA gene obtained via the Qiime2 fragment insertion plugin^65^ and molecular similarity based on Tanimoto coefficients on RDKit fingerprints. To verify the robustness of various model predictions we conducted extensive analysis, comparing the machine learning techniques listed above to various naïve models and controlling for statistical noise (see full details in Supplementary Text 1).

### Feature importance

We calculated model feature importance using the method published by Altman *et al*.^18^ using the implementation in the “ranger” package^62^. This approach allows the calculation of importance scores with statistical significance measures. To estimate the per-strain feature importance we trained the RF model with a single strain at a time and extracted the feature importance using the above method. To illustrate strain-specific differences in feature importance score, we use a principal component analysis (PCA) and demonstrated statistically significant clustering by PERMANOVA (using the “vegen” package^66^).

### Metagenomic data pre-processing

We acquired 16S rRNA amplicon sequencing data from three published studies with metagenomic longitudinal sampling obtained before and after drug treatment^38–40^, as well as from a healthy western population cohort^26^. For consistency, we processed and analyzed each dataset in a similar way. Specifically, we obtained raw fastq files from public repositories (NCBI Sequence Read Archive or European Nucleotide Archive) and processed these data using Qiime2 version 2019-1^27^. We demultiplexed the data using the Qiime2 demux plugin, applied DADA2^67^ to denoise the data, and trimmed reads in each dataset to the first position with a median quality score under 30. Since the reverse reads were of low quality, these reads were discarded. To assign ASVs to taxonomy, we trained a Naive Bayes classifier per dataset using Qiime2’s feature-classifier plugin^68^. Classifiers were trained on reads extracted from the SILVA 99-OTU database^69^, according to the specific 16S hypervariable region used in each dataset.

In each dataset, we removed samples with less than 1000 reads. We further removed rare and low abundance taxa, leaving those with abundance >0.5% in at least 0.5% of the samples. Read counts were normalized to sum to 1 within each sample, resulting in a table of relative abundances. When multiple timepoints were available, we averaged all before or after samples. Lastly, we calculated the change in relative abundance after treatment for each ASV using t-statistics.

### Predicting drug-microbe interaction landscape

To predict a rich landscape of interactions between drugs and microbes, we processed metagenomic reads from a representative healthy western population (n=90, mean age = 28)^26^. We used a single random metagenomic sample form each donor. We included in the analysis all ASV’s that appear in relative abundance above 0.5% resulting in 409 taxa in total. In parallel, we extracted from drugbank SMILES representations of all organic small molecule drugs with ATC annotations (indicating that the drug is indeed in clinical use). To maintain the credibility of predictions, we focused on small molecule drugs, removing proteins and inorganic compounds. Lastly, we trained our model on the full *in-vitro* database and predicted the new interactions.

### Finding protein of target for human drugs

We extracted known, manually curated, protein targets of drugs from the drugbank database. We then mapped those proteins to KEGG KOs via the Uniprot ID and catalogs the taxa that encode these KO in their genome. Lastly, we compared the impact score of taxa with and without the presence of the KO using the Wilcoxon rank-sum test.

### *In-vivo* prediction

As before, the impact of each drug on each microbiome member was predicted using our model. The drug features were calculated as previously described. The microbial features for each ASV were obtained from intermediate files of PICRUST2 that list KEGG KO annotation estimation^28^. Based on the full *in-vitro* data, the model predicted the sensitivity of each ASV in the sample (i.e., the impact of the given drug on that ASV). We compared the model scores for ASV with decreased relative abundance (negative t-statistics) vs. those with increased relative abundance (positive t-statistics) by two ways t-test. To further verify that the results cannot be attributed to statistical noise, we repeated this analysis on randomly shuffled data. For comparison of drug resistance between different healthy populations, we calculated the personal impact score, defined as the mean impact of drugs on each taxon and weighted by the taxa’s relative abundance.

### Associations between side effects and drug impact

We retrieved drug side effects from the SIDER database^42^. We selected side effects according to the MedDRA preferred term and all drugs with ATC annotation. Rare side effects that have been reported for less than 50 drugs were discarded. For each drug, we calculated side effect incidence frequency base on the mean frequency across all the data sources in the database and subtracted the incidence frequency of the placebo-treated groups. We grouped side effects of interest into several broader categories. Specifically, we grouped constipation, abdominal pain, diarrhea, gastrointestinal disorder, gastrointestinal pain, vomiting, nausea, abdominal discomfort, dyspepsia, flatulence, and abdominal pain upper into “Gastrointestinal side effects”. Similarly, we grouped infection, urinary tract infection, pneumonia, stomatitis, and upper respiratory tract infection into “Iinfection side effects”. We next, calculated the mean frequency across all side effects within each category. We then calculated the mean impact score of each drug across all microbes and compared this mean impact score for drugs above vs. below 1% side effect frequency.

Lastly, we explored the association between the dysbiosis pattern of drugs and their side effects. We calculated for each drug the difference in impact score between Bacteroidetes and Firmicutes. Based on this measure, we partitioned the drugs into four quartiles. Drugs in the fourth quartile were classified as firmicutes specific, drugs in the first quartile were classified as bacteroidetes specific, and drugs in the middle two quartiles were classified as not specific to any phyla.

## Supplementary Tables

**Supplementary Table 1** – A list of machine learning model features used in our framework (92 drug features and 148 microbial features)

**Supplementary Table 2** – An interaction catalogue between 2,585 drugs and 409 taxa

**Supplementary Table 3** – Possible protein targets of human-targeted drugs with statistical significance of difference in impact scores

**Supplementary Table 4** – A summary of in-vivo studies used in our analysis

## Supplementary Text

**Supplementary Text 1** – Machine learning model additional evaluation

## Notes

### Competing Interest Statement

The authors have declared no competing interest.

## References

1. Fan, Y. & Pedersen, O. Gut microbiota in human metabolic health and disease. Nat. Rev. Microbiol. 19, 55–71 (2021).

2. Zimmermann, M., Zimmermann-kogadeeva, M., Wegmann, R. & Goodman, A. L. Mapping human microbiome drug metabolism by gut bacteria and their genes. Nature doi:10.1038/s41586-019-1291-3.

3. Zimmermann, M., Zimmermann-kogadeeva, M., Wegmann, R. & Goodman, A. L. Separating host and microbiome contributions to drug pharmacokinetics and toxicity. Science (80-.). 9931, (2019).

4. Rosina Pryor, Daniel Martinez-Martinez, Leonor Quintaneiro, and F. C. The Role of the Microbiome in Drug Response. Annu. Rev. Pharmacol. Toxicol. (2020) doi:10.1007/s41974-020-00155-7.

5. Patterson, A. D. & Turnbaugh, P. J. Microbial Determinants of Biochemical Individuality and Their Impact on Toxicology and Pharmacology. Cell Metab. 20, 761–768 (2014).

6. Koppel, N., Rekdal, V. M. & Balskus, E. P. Chemical transformation of xenobiotics by the human gut microbiota. Science (80-.). 356, 1246–1257 (2017).

7. Klinken, J. B. Van et al. Integration of epidemiologic, pharmacologic, genetic and gut microbiome data in a drug–metabolite atlas. Nat. Med. 26, (2020).

8. Vich Vila, A. et al. Impact of commonly used drugs on the composition and metabolic function of the gut microbiota. Nat. Commun. 11, 1–11 (2020).

9. Jackson, M. A. et al. Gut microbiota associations with common diseases and prescription medications in a population-based cohort. Nat. Commun. 9, 1–8 (2018).

10. Forslund, K. et al. Disentangling type 2 diabetes and metformin treatment signatures in the human gut microbiota. Nature 528, 262–266 (2015).

11. Maier, L. et al. Extensive impact of non-antibiotic drugs on human gut bacteria. Nature 555, 623–628 (2018).

12. Li, L. et al. RapidAIM: a culture-and metaproteomics-based Rapid Assay of Individual Microbiome responses to drugs. Microbiome 1–16 (2020).

13. Galeano, D., Gerstein, M. & Paccanaro, A. Predicting the frequencies of drug side effects. Nat. Commun. 1–14 (2020) doi:10.1038/s41467-020-18305-y.

14. Khan, S., Hauptman, R. & Kelly, L. Engineering the Microbiome to Prevent Adverse Events: Challenges and Opportunities. Annu. Rev. Pharmacol. Toxicol. (2021).

15. Guthrie, L., Wolfson, S. & Kelly, L. The human gut chemical landscape predicts microbe-mediated biotransformation of foods and drugs. Elife 1–26 (2019).

16. Weersma, R. K., Zhernakova, A. & Fu, J. Interaction between drugs and the gut microbiome. Gut 69, 1510–1519 (2020).

17. Bajusz, D., Rácz, A. & Héberger, K. Why is Tanimoto index an appropriate choice for fingerprint-based similarity calculations ? J. Cheminform. 1–13 (2015) doi:10.1186/s13321-015-0069-3.

18. Altmann, A., Tolo, L., Sander, O. & Lengauer, T. Permutation importance: a corrected feature importance measure. Bioinformatics 26, 1340–1347 (2010).

19. Shea, R. O. & Moser, H. E. Physicochemical Properties of Antibacterial Compounds: Implications for Drug Discovery Rosemarie. J. Med. Chem. 51, (2008).

20. Brown, D. G., May-dracka, T. L., Gagnon, M. M. & Tommasi, R. Trends and Exceptions of Physical Properties on Antibacterial Activity for Gram-Positive and Gram-Negative Pathogens. J. Med. Chem. (2014).

21. Lee, J., Wood, T. K. & Lee, J. Roles of Indole as an Interspecies and Interkingdom Signaling Molecule. Trends Microbiol. 23, 707–718 (2015).

22. Aussel, L. et al. Biochimica et Biophysica Acta Biosynthesis and physiology of coenzyme Q in bacteria. BBA - Bioenerg. 1837, 1004–1011 (2014).

23. Estrada, A., Wright, D. L. & Anderson, A. C. Antibacterial Antifolates: From Development through Resistance to the Next Generation. Cold Spring Harb Perspect Med. 1–10 (2016).

24. Tommasi, R., Brown, D. G., Walkup, G. K., Manchester, J. I. & Miller, A. A. ESKAPEing the labyrinth of antibacterial discovery. Nat. Rev. Drug Discov. 14, (2015).

25. Wishart, D. S. et al. DrugBank 5.0: a major update to the DrugBank database for 2018. Nucleic Acids Res. 46, 1074–1082 (2018).

26. Poyet, M. et al. A library of human gut bacterial isolates paired with longitudinal multiomics data enables mechanistic microbiome research. Nat. Med. 25, (2019).

27. Register, F. & Services, H. Reproducible, interactive, scalable and extensible microbiome data science using QIIME 2. Nat. Biotechnol. 37, (2019).

28. Gavin M. Douglas, Vincent J. Maffei, Jesse R. Zaneveld, Svetlana N. Yurgel, James R. Brown, Christopher M. Taylor, C. H. & M. G. I. L. PICRUSt2 for prediction of metagenome functions. Nature Biotechnology vol. 38 685–688 (2020).

29. Han, Y. et al. Intestinal Dysbiosis Correlates With Sirolimus-induced Metabolic Disorders in Mice. Transplantation 105, 1017–1029 (2021).

30. Liu, Z. Z. L., Jiao, H. T. W., Xu, S. Z. Y., Mukherjee, Z. S. A. & Hu, X. Z. X. Immunosuppressive effect of the gut microbiome altered by dose tacrolimus in mice. Am J Transplant. 1646–1656 (2018) doi:10.1111/ajt.14661.

31. Li, G. et al. Gut microbiota patterns associated with somatostatin in patients undergoing pancreaticoduodenectomy: a prospective study. Cell Death Discov. (2020) doi:10.1038/s41420-020-00329-4.

32. Romualdo Barroso de Sousa, Nadim Ajami, Tanya E Keenan, Chelsea Andrews, Jessica L Pittenger, Gerburg Wulf, Laura Spring, Ian E Krop, Eric P Winer, E. A. M. and S. M. T. Fecal microbiome and association with outcomes among patients (pts) receiving eribulin (E) +/-pembrolizumab (P) for hormone receptor positive (HR+) metastatic breast cancer (MBC). Cancer Res. (2020) doi:10.1158/1538-7445.SABCS19-P3-09-16.

33. Masi, M. & Pos, K. M. influx and efflux in Gram-negative bacteria. Nat. Microbiol. (2017) doi:10.1038/nmicrobiol.2017.1.

34. Manor, O. et al. Health and disease markers correlate with gut microbiome composition across thousands of people. Nat. Commun. 1–12 doi:10.1038/s41467-020-18871-1.

35. Michael Choi, K. K. and A. K. Flavin-Dependent Thymidylate Synthase as a New Antibiotic Target. Molecules 21, 1–10 (2016).

36. He, Y., Martinez-fleites, C., Bubb, A., Gloster, T. M. & Davies, G. J. Structural insight into the mechanism of streptozotocin inhibition of O-GlcNAcase. Carbohydr. Res. 344, 627–631 (2009).

37. Hegazy, W. A. H., Khayat, M. T., Ibrahim, T. S. & Nassar, M. S. Repurposing Anti-diabetic Drugs to Cripple Quorum Sensing in Pseudomonas aeruginosa. Microorganisms (2020).

38. Seto, C. T., Jeraldo, P., Orenstein, R., Chia, N. & DiBaise, J. K. Prolonged use of a proton pump inhibitor reduces microbial diversity: Implications for Clostridium difficile susceptibility. Microbiome vol. 4 1–11 (2016).

39. Ramakrishna, C. et al. Dominant Role of the Gut Microbiota in Chemotherapy Induced Neuropathic Pain. Sci. Rep. 9, 1–16 (2019).

40. Letertre, M. P. M. et al. A Two-Way Interaction between Methotrexate and the Gut Microbiota of Male Sprague – Dawley Rats. J. Proteome Res (2020) doi:10.1021/acs.jproteome.0c00230.

41. Vangay, P. et al. Article US Immigration Westernizes the Human Gut Article US Immigration Westernizes the Human Gut Microbiome. Cell 175, 962–972.e10 (2018).

42. Kuhn, M., Letunic, I., Jensen, L. J. & Bork, P. The SIDER database of drugs and side effects. Nucleic Acids Res. 44, 1075–1079 (2016).

43. Vandeputte, D. et al. Stool consistency is strongly associated with gut microbiota richness and composition, enterotypes and bacterial growth rates. Gut 57–62 (2016) doi:10.1136/gutjnl-2015-309618.

44. Rosener, B. et al. Evolved bacterial resistance against fluoropyrimidines can lower chemotherapy impact in the Caenorhabditis elegans host. Elife 1–24 (2020).

45. Wang, Y., Bond, P. L. & Guo, J. Non-antibiotic pharmaceuticals promote the transmission of multidrug resistance plasmids through intra-and intergenera conjugation. ISME J. (2021) doi:10.1038/s41396-021-00945-7.

46. Grosheva, I. et al. High-Throughput Screen Identifies Host and Microbiota Regulators of Intestinal Barrier Function. Gastroenterology 159, 1807–1823 (2020).

47. Mahler, D. L. et al. Predicting and Manipulating Cardiac Drug Inactivation by the Human Gut Bacterium Eggerthella lenta. Science (80-.). 341, 295–299 (2013).

48. Javdan, B. et al. Personalized Mapping of Drug Metabolism by the Human Gut Microbiome. Cell 181, 1661–1679.e22 (2020).

49. Rekdal, V. M. et al. Discovery and inhibition of an interspecies gut bacterial pathway for Levodopa metabolism Discovery and inhibition of an interspecies gut bacterial pathway for Levodopa metabolism. Science (80-.). 6323, (2019).

50. Nayak, R. R. et al. Methotrexate impacts conserved pathways in diverse human gut bacteria leading to decreased host immune activation. Cell Host Microbe 29, 362–377.e11 (2021).

51. Vieira-silva, S. et al. Statin therapy is associated with lower prevalence of gut microbiota dysbiosis. Nature (2020) doi:10.1038/s41586-020-2269-x.

52. Zou, L. et al. Bacterial metabolism rescues the inhibition of intestinal drug absorption by food and drug additives. PNAS 1–10 (2020) doi:10.1073/pnas.1920483117.

53. Bhatt, A. P., Pellock, S. J., Biernat, K. A., Walton, W. G. & Wallace, B. D. Targeted inhibition of gut bacterial β-glucuronidase activity enhances anticancer drug efficacy. PNAS 117, (2020).

54. Wallace, B. D. et al. Alleviating Cancer Drug Toxicity by Inhibiting a Bacterial Enzyme. Science (80-.). 831–836 (2010).

55. Alexander, J. L. et al. Gut microbiota modulation of chemotherapy efficacy and toxicity. Nat. Rev. Gastroenterol. Hepatol. (2017) doi:10.1038/nrgastro.2017.20.

56. RDKit: Open-Source Cheminformatics Softwere. (2020).

57. Eng, A., Verster, A. J. & Borenstein, E. MetaLAFFA: a flexible, end-to-end, distributed computing-compatible metagenomic functional annotation pipeline. BMC Bioinformatics 21, 471 (2020).

58. Mukherjee, S. et al. Genomes OnLine Database (GOLD) v.8: overview and updates. Nucleic Acids Res. 49, 723–733 (2021).

59. Kanehisa, M., Sato, Y. & Morishima, K. BlastKOALA and GhostKOALA: KEGG Tools for Functional Characterization of Genome and Metagenome Sequences. J. Mol. Biol. 428, 726–731 (2016).

60. Team, R. C. R: A language and environment for statistical computing. (2020).

61. Kuhn M, W. H. Tidymodels: a collection of packages for modeling and machine learning using tidyverse principles. (2020).

62. Wright, M. N. & Ziegler, A. ranger: A Fast Implementation of Random Forests for High Dimensional Data in C ++ and R. J. Stat. Softw. 77, (2017).

63. Karatzoglou, A. & Smola, A. kernlab – An S4 Package for Kernel Methods in R. J. Stat. Softw. 11, (2004).

64. Friedman, J., Hastie, T. & Tibshirani, R. Regularization Paths for Generalized Linear Models via Coordinate Descent. J. Stat. Softw. 33, (2010).

65. Janssen, S. et al. Phylogenetic Placement of Exact Amplicon Sequences Improves Associations with Clinical Information Stefan. mSystems 3, (2018).

66. Jari Oksanen, F. Guillaume Blanchet, Michael Friendly, R. K., Pierre Legendre, Dan McGlinn, Peter R. Minchin, R. B. O., Gavin L. Simpson, Peter Solymos, M. Henry H. Stevens, E. S. & Wagner, H. vegan: Community Ecology Package. (2020).

67. Callahan, B. J. et al. DADA2: High-resolution sample inference from Illumina amplicon data. Nat. Methods 13, (2016).

68. Bokulich, N. A. et al. Optimizing taxonomic classification of marker-gene amplicon sequences with QIIME 2’s q2-feature-classifier plugin. Microbiome 1–17 (2018).

69. Pruesse, E. et al. SILVA: a comprehensive online resource for quality checked and aligned ribosomal RNA sequence data compatible with ARB. Nucleic Acids Res. 35, 7188–7196 (2007).

